# Audiograms of howling monkeys: are extreme loud calls the result of runaway selection?

**DOI:** 10.1101/539023

**Authors:** Marissa A. Ramsier, Andrew J. Cunningham, May R. Patiño, Kenneth E. Glander, Nathaniel J. Dominy

**Author notes:** Current Address: Department of Anthropology, Humboldt State University, Arcata, CA 95521, USA.

## Abstract

The eponymous vocalizations of howling monkeys (genus *Alouatta*) are associated with territorial defense and male-male competition, yet the extreme loudness of howls, which are among the loudest vocalizations of any terrestrial mammal, have yet to be fully explained. Loudness facilitates long-distance sound propagation but the effectiveness of any vocal signal depends in part on the auditory capabilities of the intended receiver, and the auditory sensitivities of howling monkeys are unknown. To better understand the evolution of loud calls, we used the auditory brainstem response (ABR) method to estimate the auditory sensitivities of *Alouatta palliata*. The mean estimated audiogram of four wild-caught adults displayed a w-shaped pattern with two regions of enhanced sensitivity centered at 0.7-1.0 and 11.3 kHz. The lower-frequency region of auditory sensitivity is pitched moderately higher than the fundamental frequencies of howling, whereas the higher-frequency region corresponds well with harmonics in an infant distress call, the *wrah-ha*. Fitness advantages from detecting infants amid low-frequency background noise, including howling, could explain the incongruity between our ABR thresholds and the fundamental frequencies of howling. Attending to infant calls is expected to enhance reproductive success within an infanticidal genus, and we suggest that the extraordinary loudness of male howling is an indirect (runaway) result of positive feedback between the selective pressures of hearing infant distress calls and deterring infanticide.

## Introduction

Howling monkeys (genus *Alouatta*) are an enduring source of fascination for vocalizing with low fundamental frequencies (300-1000 Hz [1–6]) and extreme amplitudes (70 dB re 20 at 50 m [1]), and doing so over prolonged periods, with some bouts exceeding an hour [7]. This curious combination of acoustic and behavioral traits, which Darwin described as a “dreadful concert” (p. 277 [8]), hints at substantial costs, both energetically and via increased conspicuousness to predators [9, 10]. Accordingly, researchers have long puzzled over how and why howls are emitted. Functional anatomists have examined the elaborate hyolaryngeal structures of howlers to better understand the mechanics of sound production, especially the extraordinary amplitudes and low frequencies [11–14]. Their observation that low fundamental frequencies depend on the mass and length of the vocal folds –and so vary with larynx and body size [6]– has proven instructive, leading most behavioral ecologists to view howling as an honest signal of male condition or coalition size that functions to deter conflict with conspecific rivals [15–24].

Howling is also viewed as a form of long-distance communication because loud, low-frequency sounds propagate well in forest habitats. Primate vocalizations that facilitate long-distance communication (termed “loud calls” or “long calls”) are relatively common [25] and often described as acoustic adaptations (sensu Ey and Fischer [26]). It is therefore tempting to describe howling as a quintessential acoustic adaptation [4]; indeed, humans can hear howling from a distance of 1 km [27], or possibly 5 km [28], despite high levels of background noise and attenuation. Our ability to hear distant howls can be quite striking, and some authors have called attention to the phenomenon as an exemplar of auditory stream segregation [29]. Yet, the effective range (active space) of any vocal signal depends in part on the auditory sensitivity of the intended receiver [30], and the auditory sensitivities of howling monkeys are unknown. It is this empirical void that motivated the present study of mantled howling monkeys (*Alouatta palliata*; Fig. 1A).

**Fig 1.**
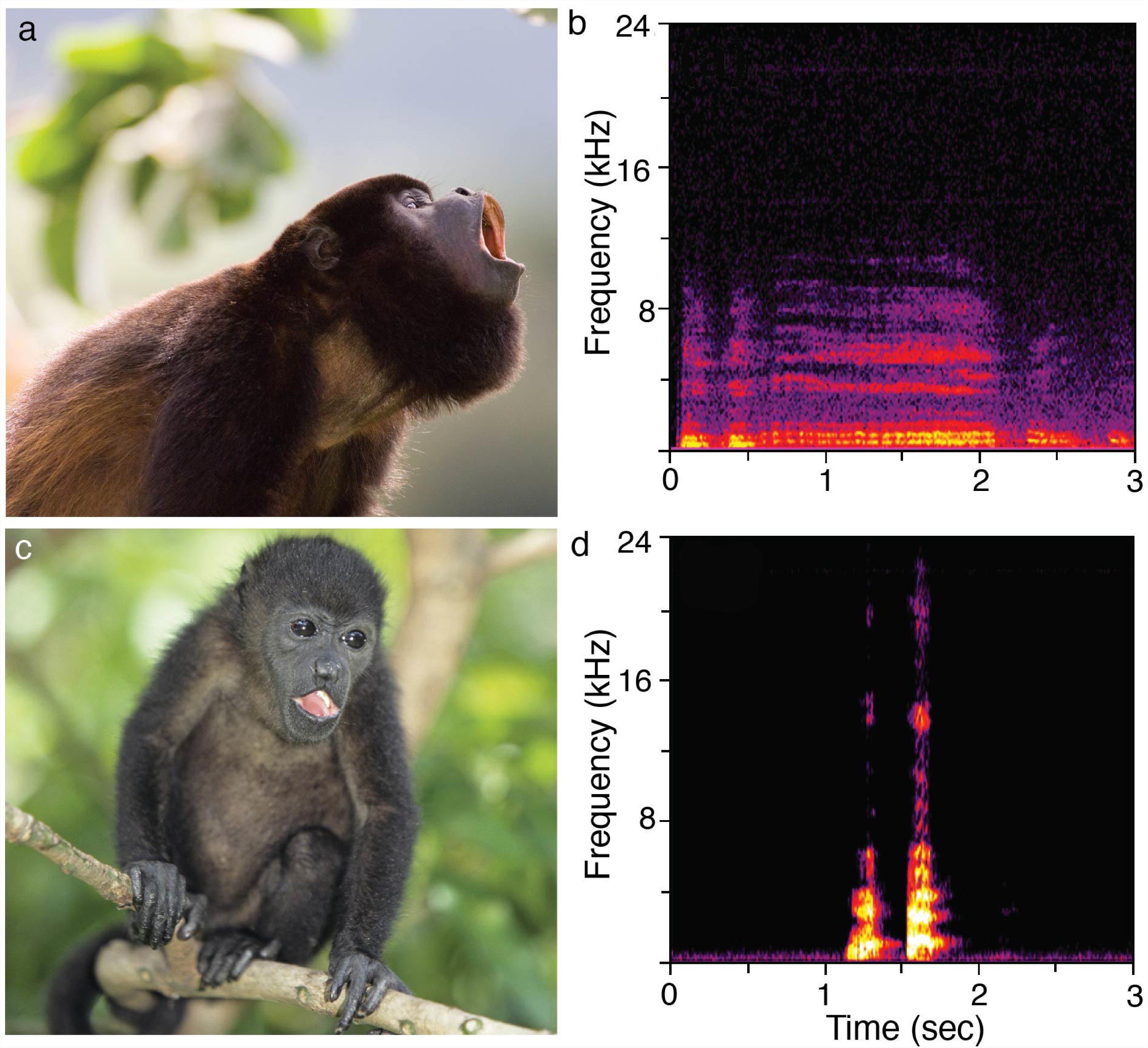
Vocal behaviors of the mantled howling monkey (*Alouatta palliata*). A: Loud calling (howling) by an adult male (photograph by David Tipling, reproduced with permission). B: Corresponding spectrogram of male howling; the audio file is available as S1 Audio. C: Unweaned infants <8 months of age emit a *wrah-ha* call when isolated from adult caregivers (photograph by Tom Brakefield, reproduced with permission). D: Corresponding spectrogram of the *wrah-ha* call; the audio file is available as S2 Audio.

The howls of *A. palliata* contain fundamental frequencies between 300 and 500 Hz [2–5] (Fig. 1B), and it follows that adult receivers will have commensurate hearing sensitivities to optimize the range of this vocal signal, a concept termed frequency matching. Frequency matching can evolve because the auditory system has shaped the evolution of important vocal signals (sensory drive [31]) or because important vocal signals have shaped the evolution of the auditory system (social drive [32]). At the same time, the unweaned infants of *A. palliata* are known to produce a conspicuous, high-frequency isolation/distress call when separated from caregivers or during weaning (Fig. 1C). This call is termed the *wrah-ha* [7] (Fig. 1D; cf. call type 5 of Carpenter [33] and whimpers of Altmann [34] for further context). The fitness benefits of attending to the *wrah-ha* could have imposed constraints on the hearing of adults, biasing their auditory sensitivity toward higher frequencies. This concept is termed receiver bias [31] and the effect on adults is potentially greatest when the risk of infanticide is high. Reports of infanticide in *Alouatta* [35–39] (including *A. palliata* [40, 41]) distinguish howling monkeys from all other neotropical primates, and it is plausible that the auditory sensitivities of adults have been shaped in part by the vocalizations of imperiled infants.

We highlight these calls of *A. palliata* because key frequencies are expected to fall at opposite ends of the audible range of adult receivers, and because there is evidence for evolutionary trade-offs in primate hearing sensitivity [42]; i.e., adults are unlikely to be equally sensitive to extremely low and high frequencies. To explore whether the auditory sensitivities of adult receivers are better matched to the howling of adult males or the *wrah-ha* of infants, we collected field recordings of each call and used the auditory brainstem response (ABR) method to generate audiograms from wild-caught individuals. The ABR method records neural responses to acoustic stimuli, and it has been used to estimate audiograms from a wide range of nonhuman primates in captive and wild settings [32, 43–46]. The main advantages of the ABR method are its portability and efficiency, and the compatibility of ABR-derived audiograms with behavioral audiograms in overall shape and at least two important parameters: (i) the frequency of best sensitivity and (ii) the high-frequency limit (the highest frequency that can be detected at at 60 dB SPL) [43]. A disadvantage of the method is that thresholds ≤2 kHz are often elevated compared to behavioral thresholds, and therefore represent maximum (“at least this sensitive”) estimates rather than absolute threshold values [43].

## Materials and methods

### Ethics statement

This research complied with protocols approved by the Ministerio de Ambiente y Energía (MINAE) of Costa Rica (permit no. 04301) and the Institutional Animal Care and Use Committees of Humboldt State University (approval no. 11/12.A.105-A), Dartmouth College (approval no. 10-11-0), Duke University (approval nos. A003-0801 and A325-10-12) and the University of California, Santa Cruz (approval nos. Domin0711 and Domin0810).

### Animal capture and study conditions

We tested mantled howling monkeys (*Alouatta palliata*) inhabiting the seasonal deciduous forests of La Pacifica, Costa Rica. The life histories of individual monkeys at this site are well known due to 40 years of mark and recapture studies [47, 48]. We captured animals with Pneu-Darts (Williamsport, PA, USA) loaded with Telazol (25 mg/kg) [49, 50] and used supplemental doses of Telazol (2 mg/kg) or medetomidine (45/kg IM) to sustain anesthesia during the testing period (≈1 hr). We used these anesthetics to minimize subject stress and interference from myogenic noise; neither is known to have significant effects on ABR amplitudes [43]. We monitored the vital parameters of each animal and used electric heating pads to maintain body temperatures as needed. At the conclusion of ABR testing, we reversed (45/kg atipanezole IM), monitored (for several hours), and released each animal unharmed within its home range.

We tested four individuals (subjects H, I, J, and K) in July 2008 and March 2011 (Table 1). Data from seven additional animals (subjects A-F) were collected in February 2008 and reported in an unpublished PhD thesis [51]; however, these data are excluded here because we have since determined that high electroencephalographic (EEG) background noise and stimulus artifacts adversely affected the ABR.

**Table 1.** Identifications and attributes of each study animal

| Animal identifications | Sex | Age (years) | Mass (kg) |
| --- | --- | --- | --- |
| H (“Ruapehu”) | male | young adult | 6.0 |
| I (“Riley”) | male | $\approx 8.0$ | 7.0 |
| J (“May”) | female | 4.5 | 4.5 |
| K (“Almeda”) | female | $\approx 5.5$ | 6.3 |

Subjects H, I, J, and K were put into a supine position with the left ear facing the stimulus. Subject H was tested in a 16-m^2^ room, in which ambient acoustic noise fluctuated from 15-30 dB SPL across the frequency range tested (1.0-32.0 kHz). Subjects I, J, and K were put into an 80.0 x 67.3 x 63.5-cm sound-attenuating cubicle (model 83017DDW, Lafayette Instrument Co., Lafayette, IN, USA) constructed with thick (19 mm) moderately expanded PVC foam panels lined with thick (53 mm) painted acoustical foam (SONEX, Minneapolis, MN, USA). Ambient acoustic noise in the cubicle fluctuated from 0-20 dB SPL (1.0-64.0 kHz) and intermittently reached as high as 29 dB at 0.7 kHz and 56 dB at 0.5 kHz. For each monkey and tested frequency, the EEG background noise was consistently low and the response amplitude decreased steadily with decreasing stimulus levels (average r2: 0.83), indicating minimal masking from ambient noise. Even still, the thresholds for frequencies <1 kHz should be considered preliminary.

### ABR stimuli and recording

ABR stimulus generation and response acquisition was accomplished with EVREST software [52] operating on a laptop PC equipped with a data acquisition card (NI-USB 6251, National Instruments, Austin, TX, USA). The stimuli were tone pips (2-cycle linear rise/fall, 1-cycle plateau) generated digitally, converted to analog (500 kHz, 16-bit resolution), bandpass filtered from 0.02 to 200 kHz (3-B series, Krohn-Hite, Brockton, MA, USA; 24 dB/octave rolloff, Butterworth), attenuated (PA5, Tucker-Davis, Alachua, FL, USA), and delivered through a free field via a magnetic speaker (FF1, Tucker-Davis; frequency response <1-50 kHz) or an electrostatic speaker (ES1, Tucker-Davis; frequency response 4-110 kHz) positioned 10 cm from the left ear. The test frequencies were half octaves from 1.0-32.0 kHz (subject H) or 0.5-32.0 kHz (subjects I, J, and K), delivered in steps of 10 or 5 dB starting ≈30 dB above the estimated threshold and decreasing to at least 5 dB below the level where the response fell to residual EEG background noise (BN) level and was undetectable. Each frequency/stimulus condition was delivered for 1024-2048 repetitions at a rate of 39.1 sec^-1^.

The levels of the tone-pip stimuli were calibrated before the testing of the first subject, after the last subject, and periodically throughout each field season. While in the field, 50-100 ms pure tones were produced with equal system output voltage to the tone-pip stimuli, and recorded via a free-field 0.5-inch condenser microphone (MKH 800, Sennheiser, Old Lyme, CT, USA; frequency response 0.03-50 kHz, 0°) connected to a PC running Raven Pro v. 1.3 (Cornell Laboratory of Ornithology, Ithaca, NY, USA). Within Raven, we calculated dB SPL (re 20Pa) with the absolute level calibrated at 1 kHz via a Type-1 sound level meter (840015, Sper Scientific, Scottsdale, AZ; C frequency weighting). Under subsequent lab conditions, we accounted for the nonlinear frequency response of the MKH 800 microphone by recording the same stimuli with the MKH 800 and a 0.25-inch free-field microphone (Brüel and Kajer 4939, Nærum, Denmark; frequency response 0.004-100 kHz, 0°) calibrated with a class 1 sound calibrator (Brüel and Kajer, Nærum, Denmark). Equal system output peak-peak voltage of the pure tones and tone pips was also verified with a 32-bit digital oscilloscope (DSO Nano, Seeed Studio, Shenzhen, China) (vis-à-vis the peSPL method of Laukli and Burkard [53]).

We recorded the ABR via 28-gauge subdermal needle electrodes (F-E3, Grass Instruments, West Warwick, RI, USA) inserted in the skin over the cranial vertex (positive) and the ipsilateral mastoid (reference); a ground was placed over the contralateral mastoid. Signals were recorded via a biopotential amplifier (P511, Grass Instruments), amplified (x 105), filtered (0.03-3 kHz bandpass, 0.06 kHz notch), digitized (10 kHz, 16-bit resolution), input into EVREST in 20 ms epochs (reject level 12), and digitally filtered offline.

### Data analysis

For each subject, we used linear regression to determine the threshold (quietest detectable level) for each presented frequency [43]. We plotted the response amplitude as a function of stimulus level – generally, all responses above average BN level were included. Next, we used the line of best fit (mean r2: 0.83) to determine the level at which the response reached a threshold criterion of 50 nV (average BN + 40 nV - after Ramsier et al. [44]); at this criterion, the response was consistently above average random fluctuation in BN (S1 Fig).

To calculate the mean estimated audiogram for the species, we estimated some individual thresholds in order to avoid biasing the average and distorting the overall shape of the audiogram. Some individual variation in absolute sensitivities is expected (or at least the measurement of such; i.e., an up or down shift of the audiogram), but the overall shape of an audiogram varies little at the species level [32, 43, 44]. Thus, we estimated missing threshold values by using the average rise/fall from adjacent frequencies in other individuals. We then computed the frequency of best sensitivity and the high-frequency limit; we focused our analysis on these parameters because they can be estimated reliably with the ABR method [43].

### Field recordings and preliminary playback experiments

During the course of our study, we opportunistically recorded howls using a free-field 0.5-inch condenser microphone (MKH 800, Sennheiser, Old Lyme, CT, USA; frequency response 0.03-50 kHz, 0°) connected to a solid-state recorder (Marantz PMD-671, sampling rate 96 kHz). We also recorded infants emitting *wrah-ha* calls, which we observed under three conditions: (i) when separated from their mother or natal group; (ii) during human handling; or (iii) when seeking attention from unresponsive adults. Audio files of howling and the *wrah-ha* call are available as S1 Audio and S2 Audio, respectively. Spectrograms of each vocalization are illustrated in Fig. 1B and Fig. 1D, respectively.

To explore the function and potential fitness costs of the infant *wrah-ha* call, we performed pilot playback studies in the home ranges of howlers at two field sites in Costa Rica: La Pacifica (seasonal dry forest) and El Zota (lowland rain forest). The calls were broadcast from a lifelike rubber model intended to represent an infant on the verge of weaning, at ≈8 months of age [54] (Animal Makers Inc., Moorpark, CA, USA; S2 Fig). In most conditions, a loudspeaker (Micro Cube, Roland Corp., Hamamatsu, Japan) was positioned next to the model. In one condition, a magnetic speaker (FF1, Tucker-Davis) was embedded in the abdominal cavity of the rubber model. The speaker and model were positioned on tree branches ≈2 m from the ground.

## Results and Discussion

### ABR thresholds of *Alouatta palliata*

We captured four adult monkeys (Table 1) and recorded ABR thresholds from each (S1 Table). We observed considerable inter-individual variation in our sample, especially at lower frequencies. For example, the thresholds of Subjects K and I differed from those of H and J by more than 25 dB at 0.5 kHz, a large difference that persisted up to ≈1.5 kHz. Despite this level of variation, we calculated a mean estimated audiogram for comparative purposes. It displays a w-shaped pattern with two frequency regions of enhanced auditory sensitivity (frequencies of best sensitivity) separated by a mid-frequency ‘dip’ of decreased sensitivity (Fig. 2). The mean threshold at 0.5 kHz is 5.8 dB less sensitive than at 0.7 kHz, which suggests that 0.7-1.0 kHz is the lower-frequency region of enhanced sensitivity for *A. palliata*. However, a difference of 5.8 dB is trivial given the magnitude of variation between individuals at 0.5 kHz, and it is plausible that the lower-frequency region of enhanced sensitivity extends below 500 Hz. This uncertainty is a source of disappointment given our interest in frequencies between 300 and 500 Hz. The high-frequency area of auditory sensitivity was centered between 8.0-11.3 kHz. The highest frequency limit (the highest frequency detectable at 60 dB) was estimated at 21.7 kHz; however, we consider this result preliminary, as the higher value of Subject J suggests that a larger sample could yield a higher high-frequency limit.

**Fig 2.**
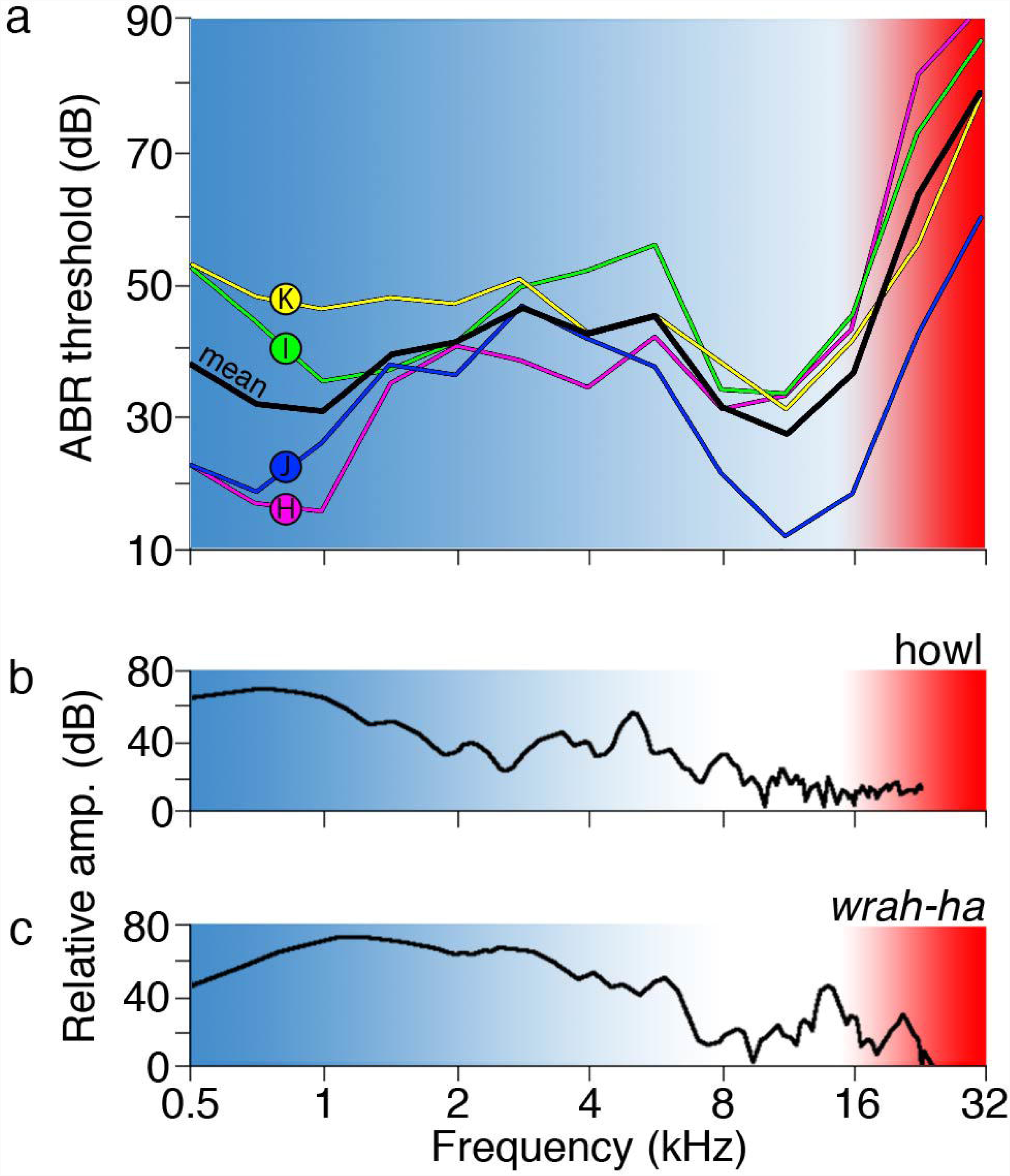
Estimated audiograms of (*Alouatta palliata*). A: The mean audiogram is shown in black together with the individual audiograms of subjects H (magenta), I (green), J (purple), and K (yellow). The red shaded region indicates ultrasound. B: Spectrogram time slice of a typical male howl. C: Spectrogram time slice of a typical infant *wrah-ha* vocalization.

The limitations of our study are threefold. First, the sample size is quite small, which reflects the challenges of capturing wild animals and testing them under field conditions. Second, we were unable to reliably estimate ABR thresholds below 500 Hz. Third, we observed substantial inter-individual variation, which introduces considerable uncertainty when attempting to estimate a mean audiogram for comparative purposes. Our data are therefore incomplete and highly variable, yet the shape of our mean estimated audiogram agrees well with those of other platyrrhine (New World) monkey species [55]. For example, the lower-frequency region of enhanced sensitivity reported here (0.7-1 kHz) is lower than or comparable with those of *Aotus trivirgatus* (2 kHz [56]), *Callithrix jacchus* (1 kHz [57]), *Saimiri sciureus* (2 kHz [58]), and *Sapajus* (*Cebus*) *apella* (4 kHz [46]). Similarly, the higher-frequency area of auditory sensitivity (8.0-11.3 kHz) resembles those of *A. trivirgatus* (10 kHz [56]), *C. jacchus* (7 kHz [57]), *S. sciureus* (12 kHz [58]), and *S. apella* (8-16 kHz [46]). This level of congruence suggests that the audiograms of platyrrhine monkeys are relatively conserved across species.

Enhanced auditory sensitivity to higher frequencies is hardly surprising for *A. trivirgatus, C. jacchus*, and *S. sciureus* on the basis of their small body sizes (they are each <1 kg). In contrast, the mass of *A. palliata* is more than sixfold greater (Table 1) and predicted on theoretical grounds to have diminished sensitivity to higher frequencies [59]. In accordance with this view, we found that the ABR threshholds of *A. palliata* increased abruptly at frequencies >16 kHz, resulting in a preliminary high-frequency limit of 21.7 kHz. What is surprising, however, is that estimated auditory sensitivity is quite good for frequencies ranging from ≈8 kHz to the lower margin of ultrasound (Fig. 2), where it diminished steeply.

### Frequency matching with vocal signals

Audiograms with a w-shaped pattern may underlie discrete ‘channels’ of communication: a lower-frequency channel used by adults and a higher-frequency channel used by infants [60]. Our findings are compatible with this view, as they point tentatively to a low-frequency area of enhanced sensitivity (0.7-1.0 kHz) that is reasonably compatible with the dominant energy of conspecific howls (≈1.4 kHz, Fig. 2B). This result suggests that *A. palliata* can hear its own howls relatively well, especially given that our low-frequency ABR thresholds are conservative estimates of auditory sensitivity. Even still, it is perhaps intriguing that the low-frequency area of auditory sensitivity is pitched moderately higher than the fundamental frequencies of howling between 300 and 500 Hz [2–5]. This incongruity must be viewed with caution, however, as we could not reliably record ABR thresholds below 500 Hz. The high-frequency area of auditory sensitivity (8.0-11.3 kHz) is shifted away from the higher-frequency components of howls and toward the *wrah-ha* with its high-frequency harmonics at ≈10 and ≈21 kHz (Fig. 2C). A comparable level of frequency matching can be observed in *Aotus trivirgatus* and *Callithrix jacchus*. In these monkeys, the frequencies of greatest auditory sensitivity are closely matched to the highest-frequency calls in their vocal repertoires —the squeals, squeaks, and trills of infants and juveniles [61, 62].

The harmonic at ≈21 kHz (Fig. 2C) is a new finding, in part because it lies beyond the frequency ranges of the recording equipment used in previous studies. It is probably the consequence of a short supralarygeal vocal tract; however, there are at least two advantages to producing vocal signals at these high frequencies. First, they should be conspicuous amid bouts of howling and the low-frequency ambient noise of tropical forests [63]. Second, the high-frequency components should clearly identify the caller as an infant and improve adult response time in the presence of predators or infanticidal males. Such mutually compatible advantages could have exerted a strong selective pressure on both the signaler and receiver, resulting in adult auditory sensitivities that are biased (upshifted) toward the higher frequencies emitted by infants. To explore this premise further, we performed preliminary playback experiments with a lifelike infant model, but the results were equivocal—the *wrah-ha* elicited furtive behaviors from adults, but no physical interactions (see S2 Fig).

### Extreme loudness of howling: is it an example of runaway selection?

Available evidence points to a “sound window” of least attenuation in the mid-canopy layers of tropical forests [64, 65]. In one oft-cited experiment, optimal sound propagation at a height of 15 m occurred between 200 and 300 Hz [64], a frequency range that lies just below the fundamental frequencies of howling by *A. palliata* (300-500 Hz; Fig. 1B) and the low-frequency region of enhanced auditory sensitivity reported here (0.7-1.0 kHz; Fig. 2A). This latter result is, admittedly, disappointing –we stress that our study faced technical limitations, and we cannot rule out a lower low-frequency region of enhanced auditory sensitivity– but it is also tantalizing, as it hints at incongruities between howling and hearing, a possibility that invites replication and warrants further discussion, albeit with due caution. If our results are verified, then an acoustic mismatch between the fundamental frequencies of howling and the low-frequency region of enhanced auditory sensitivity can be interpreted as evidence of receiver bias and a lower-frequency limit on acoustic communication [66]. Such an interpretation is alluring because it begins to explain why howls are so loud —the extreme amplitude is a compensatory mechanism to increase the range of the vocal signal, which holds particular importance for male fitness.

For dominant males, the amplitude of howling could be an honest or exaggerated signal of male body mass [67, 68]. Regardless, male body mass is a strong predictor of takeover success by invader or usurper (resident) males [47], and louder howling probably functions to repel rival males and extend the reproductive advantages of dominant males [9, 15]. Yet, delayed takeovers must carry reproductive opportunity costs for subordinate males, increasing their incentive to commit infanticide when takeovers do occur (Fig. 3). Infanticide by resident or returning males has been reported [38, 48], but the probability of paternity remains low [69, 70]. Thus, selection for louder howling is expected to attend an ever greater need to resolve and respond to the distress calls of infants. The resulting receiver bias toward detecting high frequencies is then expected to reinforce constraints on auditory sensitivity to lower frequencies and, in turn, favor louder howling instead of lower fundamental frequencies for greater signal transmission [6] (Fig. 3). Such positive feedback is reminiscent of runaway selection.

**Fig 3.**
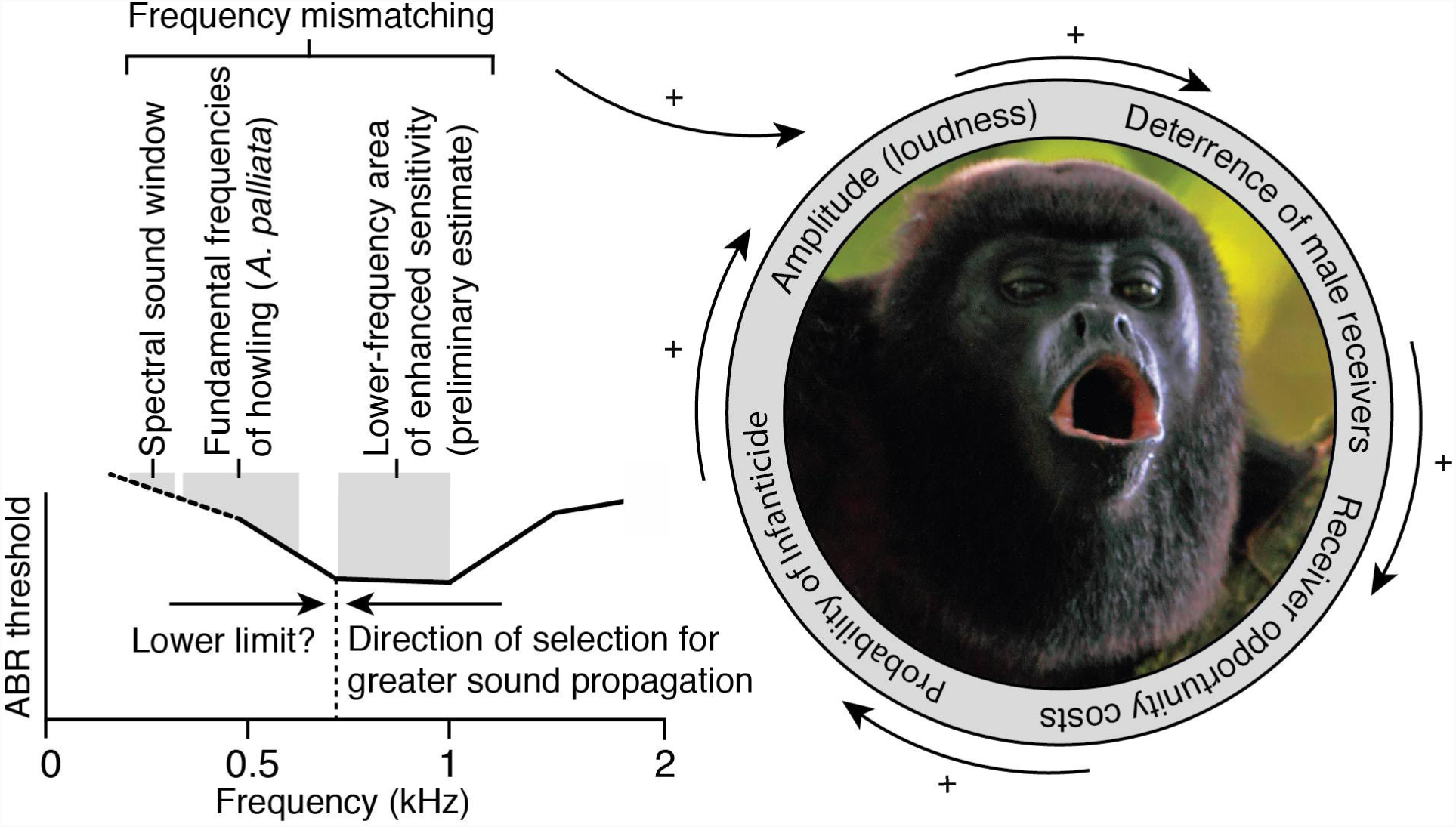
Integrated model of runaway selection. Our results suggest a lower-frequency region of enhanced auditory sensitivity that is suboptimally matched to the “sound window” of least attenuation and the fundamental frequencies of conspecific howls. This incongruity raises the possibility of a lower limit on the lower-frequency region of enhanced auditory sensitivity, perhaps because adult sensitivity is biased toward the high-frequency vocalizations of infants. This trade-off is predicted to favor louder, rather than lower-frequency howling for greater signal propagation. Louder howling, in turn, is expected to benefit signaler males by deterring receiver males and prolonging access to group females. At the same time, lone receiver males will face increasing reproductive opportunity costs that will increase the likelihood of committing infanticide during group takeover. Either male could gain fitness advantages by detecting infants in distress. Eventually, the receiver male becomes the signaler male, closing the positive feedback loop that drives ever louder howling. Thus, we hypothesize that the extraordinary loudness of howling is an indirect (runaway) result of suboptimal frequency matching and the fitness advantages of repelling lone receiver males. Photograph by Christian Ziegler, reproduced with permission.

The runaway selection hypothesis was conceived to explain the extreme elaboration of male secondary sexual characteristics [71]. It begins with a preexisting female preference for a male trait that drives further evolution of both the preferred trait and the female preference for it. In studies of animal acoustic communication, support for the runaway selection hypothesis is sparse [72, 73], and it is questionable whether any mating signal has evolved via this mechanism [74]. As Ryan and Kime [73] put it, “there are no good examples of studies in acoustic communication supporting the runaway hypothesis” (p. 253). However, runaway selection need not rely solely on female preferences. Several authors have invoked runaway selection in the context of long-distance calls related to territorial defense [75] or loud calls related to male-male competition [76], behaviors that have long factored into explanations for the extreme loudness of male howling. Our preliminary study of ABR thresholds in *A. palliata* reinforces this view and we hypothesize that the extraordinary loudness of howling is an indirect (runaway) result of positive feedback between the selective pressures of hearing infant distress calls and deterring infanticide.

## Conclusion

*Extraordinary claims require extraordinary evidence* is a popular aphorism that weighs on our minds. The present report makes no extraordinary claims; rather, our philosophy is to report admittedly incomplete and variable findings from an extraordinary animal. Research is a privilege and we view data-sharing as a moral responsibility. At the same time, the data are tantalizing, leading us to hypothesize that the extraordinary loudness of male howling is a runaway result of positive feedback between the selective pressures of hearing infant distress calls and deterring infanticide. We offer this hypothesis not as a conclusion, but as an open invitation to future research.

## Acknowledgments

For practical and technical support, we thank the staff at La Pacifica and the following individuals: M. Bezanson, D. Brenneis, D. Carstens, B.E. Crowley, J.J. Finneran, A. Galloway, M.E. Glenn, C.A. Juarez, D. Ramsier, J.M. Ratcliffe, D. Riggs, F. Spoor, S. Streiber, F. Suarez, M.F. Teaford, C. Vinyard, and S. Williams. Funding was received from the Department of Anthropology, University of California, Santa Cruz (to M.A.R.); the Max Plank Society (to M.A.R.); the National Science Foundation (grant numbers BCS-0720028, BCS-0720025, and BCS-0507074 to K.E.G.); and the David and Lucile Packard Foundation (Fellowship in Science and Engineering 2007-31754 to N.J.D.).

## Supporting information

**S1 Audio. Howling of *Alouatta palliata***. Audio recording of howling (or roaring, or loud calling) by an adult male.

**S2 Audio. *wrah-ha* of *Alouatta palliata***. Audio recording of the *wrah-ha* emitted from an infant male.

**S1 Fig. Typical ABR waveforms evoked with 11.3 kHz stimuli**. A: waveforms for each subject. B: waveform series demonstrating decreasing ABR amplitude with decreasing stimulus level.

**S2 Fig. Model infant**. Field playback experiments of the *wrah-ha* were performed in tandem with a rubber model (Animal Makers Inc., Moorpark, CA, USA). In most conditions, a loudspeaker (Micro Cube, Roland Corp., Hamamatsu, Japan) was positioned next to the model. In one condition, a magnetic speaker (FF1, Tucker-Davis) was embedded in the abdominal cavity of the rubber model. The speaker and model were positioned on tree branches ≈2 m from the ground. On 7 of 11 occasions, the vocalization quickly (within 5 min) attracted lone males or mixed groups. Animals descended within a few meters of the model, but never contacted it. In all cases, adult males and/or females demonstrated heightened vigilance and sustained howling or barking over a span of 5-20 min. On one occasion the call attracted a female black-handed spider monkey (*Ateles geoffroyi)*. Such results are equivocal, but they suggest that adults are motivated to approach distressed infants. Our failure to elicit physical interaction with the model could be attributed to the deterrent presence of human observers or the unnatural appearance and vocal ‘behavior’ of the model. Normative infant behavior predicts the suppression of distress calls in the vicinity of potentially infanticidal males. Another factor was the highly irregular social context. Unweaned infants of *A. palliata* are seldom isolated; they maintain close proximity to their mothers (< 3 m [54]) and other members of their natal group (1-5 m [77]). Our model, though lifelike from a distance, may have appeared too unnatural to elicit further response. Photograph by Andrew J. Cunningham.

**S1 Table. ABR thresholds.**

## References

1. Sekulic R. Male relationships and infant deaths in red howler monkeys (*Alouatta seniculus*). Zeitschrift für Tierpsychologie. 1983;61(3):185–202.

2. Eisenberg JF. Communication mechanisms and social integration in the black spider monkey, *Ateles fusciceps robustus*, and related species. Smithsonian Contributions to Zoology. 1976;213:1–108.

3. Thorington RW, Ruiz JC, Eisenberg JF. A study of a black howling monkey (*Alouatta caraya*) population in northern Argentina. American Journal of Primatology. 1984;6(4):357–366.

4. Whitehead JM. Vox Alouattinae: a preliminary survey of the acoustic characteristics of long-distance calls of howling monkeys. International Journal of Primatology. 1995;16(2):121–144.

5. Bergman TJ, Cortés-Ortiz L, Dias PAD, Ho L, Adams D, Canales-Espinosa D, et al. Striking differences in the loud calls of howler monkey sister species (*Alouatta pigra* and *A. palliata*). American Journal of Primatology. 2016;78(7):755–766.

6. Dunn JC, Halenar LB, Davies TG, Cristobal-Azkarate J, Reby D, Sykes D, et al. Evolutionary trade-off between vocal tract and testes dimensions in howler monkeys. Current Biology. 2015;25(21):2839–2844.

7. Baldwin JD, Baldwin JI. Vocalizations of howler monkeys (*Alouatta palliata*) in southwestern Panama. Folia Primatologica. 1976;26(2):81–108.

8. Darwin C. The descent of man and selection in relation to sex. London: John Murray; 1871.

9. Kitchen DM. Alpha male black howler monkey responses to loud calls: effect of numeric odds, male companion behaviour and reproductive investment. Animal Behaviour. 2004;67(1):125–139.

10. Stoddard PK, Salazar VL. Energetic cost of communication. Journal of Experimental Biology. 2011;214(2):200–205.

11. Kelemen G, Sade J. The vocal organ of the howling monkey (*Alouatta palliata*). Journal of Morphology. 1960;107(2):123–140.

12. Schön MA. On the mechanism of modulating the volume of the voice in howling monkeys. Acta Oto-Laryngologica. 1970;70(5-6):443–447.

13. Schön MA. The anatomy of the resonating mechanism in howling monkeys. Folia Primatologica. 1971;15(1-2):117–132.

14. schön Ybarra MA. Loud calls of adult male red howling monkeys (*Alouatta seniculus*). Folia Primatologica. 1986;47(4):204–216.

15. Sekulic R. The function of howling in red howler monkeys (*Alouatta seniculus*). Behaviour. 1982;81(1):38–54.

16. Chiarello AG. Role of loud calls in brown howlers, *Alouatta fusca*. American Journal of Primatology. 1995;36(3):213–222.

17. Kitchen DM, Horwich RH, James RA. Subordinate male black howler monkey (*Alouatta pigra*) responses to loud calls: experimental evidence for the effects of intra-group male relationships and age. Behaviour. 2004;141(6):703–723.

18. Kitchen DM. Experimental test of female black howler monkey (*Alouatta pigra*) responses to loud calls from potentially infanticidal males: effects of numeric odds, vulnerable offspring, and companion behavior. American Journal of Physical Anthropology. 2006;131(1):73–83.

19. da Cunha RGT, Byrne RW. Roars of black howler monkeys (*Alouatta caraya*): evidence for a function in inter-group spacing. Behaviour. 2006;143(10):1169–1199.

20. Holzmann I, Agostini I, Di Bitetti M. Roaring behavior of two syntopic howler species (*Alouatta caraya* and *A. guariba clamitans*): evidence supports the mate defense hypothesis. International Journal of Primatology. 2012;33(2):338–355.

21. Hopkins ME. Relative dominance and resource availability mediate mantled howler (*Alouatta palliata*) spatial responses to neighbors’ loud calls. International Journal of Primatology. 2013;34(5):1032–1054.

22. Van Belle S, Estrada A, Garber PA. Spatial and diurnal distribution of loud calling in black howlers (*Alouatta pigra*). International Journal of Primatology. 2013;34(6):1209–1224.

23. Van Belle S, Estrada A, Garber PA. The function of loud calls in black howler monkeys (*Alouatta pigra*): food, mate, or infant defense? American Journal of Primatology;76(12):1196–1206.

24. Kitchen DM, da Cunha RGT, Holzmann I, de Oliveira DAG. In: Kowalewski MM, Garber PA, Cortés-Ortiz L, Urbani B, Youlatos D, editors. Function of loud calls in howler monkeys; 2015. p. 369–399.

25. Mitani J, Stuht J. The evolution of nonhuman primate loud calls: acoustic adaptation for long-distance transmission. Primates. 1998;39(2):171–182.

26. Ey E, Fischer J. The “Acoustic Adaptation Hypothesis” – a review of the evidence from birds, anurans, and mammals. Bioacoustics. 2009;19:21–48.

27. Horwich RH, Gebhard K. Roaring rhythms in black howler monkeys (*Alouatta pigra*) of Belize. Primates. 1983;24(2):290–296.

28. Anonymous. Guinness World Records 2008. New York: Guinness World Records Limited; 2007.

29. Ghazanfar AA, Santos LR. Primate brains in the wild: the sensory bases for social interactions. Nature Reviews Neuroscience. 2004;5(8):603–616.

30. Bradbury JW, Vehrencamp SL. Principles of animal communication, Second Edition. Sunderland: Sinauer; 2011.

31. Endler JA, Basolo AL. Sensory ecology, receiver biases and sexual selection. Trends in Ecology & Evolution. 1998;13(10):415–420.

32. Ramsier MA, Cunningham AJ, Finneran JJ, Dominy NJ. Social drive and the evolution of primate hearing. Philosophical Transactions of the Royal Society B: Biological Sciences. 2012;367(1597):1860–1868.

33. Carpenter CR. A field study of the behavior and social relations of howling monkeys. Comparative Psychology Monographs. 1934;10:1–116.

34. Altmann SA. Field observations on a howling monkey society. Journal of Mammalogy. 1959;40(3):317–330.

35. Agoramoorthy G, Rudran R. Infanticide by adult and subadult males in free-ranging red howler monkeys, Alouatta seniculus, in Venezuela. Ethology. 1995;99(1-2):75–88.

36. Crockett CM, Janson CH. In: van Schaik CP, Janson CH, editors. Infanticide in red howlers: female group size, male membership, and a possible link to folivory. Cambridge: Cambridge University Press; 2000. p. 75–98.

37. Crockett CM. In: Jones CB, editor. Re-evaluating the sexual selection hypothesis for infanticide by Alouatta males. Norman: American Society of Primatologists; 2003. p. 327–365.

38. Knopff KH, Knopff ARA, Pavelka MSM. Observed case of infanticide committed by a resident male central American black howler monkey (*Alouatta pigra*). American Journal of Primatology. 2004;63(4):239–244.

39. Van Belle S, Kulp AE, Thiessen-Bock R, Garcia M, Estrada A. Observed infanticides following a male immigration event in black howler monkeys, Alouatta pigra, at Palenque National Park, Mexico. Primates. 2010;51(4):279–284.

40. Clarke MR. Infant-killing and infant disappearance following male takeovers in a group of free-ranging howling monkeys (*Alouatta palliata*) in Costa Rica. American Journal of Primatology. 1983;5(3):241–247.

41. Clarke MR, Zucker EL, Glander KE. Group takeover by a natal male howling monkey (*Alouatta palliata*) and associated disappearance and injuries of immatures. Primates. 1994;35(4):435–442.

42. Coleman MN, Colbert MW. Correlations between auditory structures and hearing sensitivity in non-human primates. Journal of Morphology;271(5):511–532.

43. Ramsier MA, Dominy NJ. A comparison of auditory brainstem responses and behavioral estimates of hearing sensitivity in *Lemur catta* and *Nycticebus coucang*. American Journal of Primatology. 2010;72(3):217–233.

44. Ramsier MA, Cunningham AJ, Moritz GL, Finneran JJ, Williams CV, Ong PS, et al. Primate communication in the pure ultrasound. Biology Letters. 2012;8(4):508–511.

45. Schopf C, Zimmermann E, Tünsmeyer J, Kästner SBR, Hubka P, Kral A. Hearing and age-related changes in the gray mouse lemur. Journal of the Association Research in Otolaryngology. 2014;15(6):993–1005.

46. Ramsier MA, Vinyard CJ, Dominy NJ. Auditory sensitivity of the tufted capuchin (*Sapajus apella*), a test of allometric predictions. Journal of the Acoustical Society of America. 2017;141(6):4822–4831.

47. Glander KE. Dispersal patterns in Costa Rican mantled howling monkeys. International Journal of Primatology. 1992;13(4):415–436.

48. Clarke MR, Glander KE. Secondary transfer of adult mantled howlers (*Alouatta palliata*) on Hacienda La Pacifica, Costa Rica: 1975–2009. Primates. 2010;51(3):241–249.

49. Glander KE, Fedigan LM, Fedigan L, Chapman C. Field methods for capture and measurement of three monkey species in Costa Rica. Folia Primatologica. 1991;57(2):70–82.

50. Glander KE, Nisbett RA. Community structure and species density in tropical dry forest associations at Hacienda La Pacífica in Guanacaste province, Costa Rica. Brenesia. 1996;45-46:113–142.

51. Ramsier MA. The evolutionary ecology of primate auditory sensitivity [PhD thesis]; University of California, Santa Cruz, USA, 2010.

52. Finneran JJ. Evoked response study tool: a portable, rugged system for single and multiple auditory evoked potential measurements. Journal of the Acoustical Society of America. 2009;126(1):491–500.

53. Laukli E, Burkard R. Calibration/standardization of short-duration stimuli. Seminars in Hearing. 2015;36(1):3–10.

54. Clarke MR. Behavioral development and socialization of infants in a free-ranging group of howling monkeys (*Alouatta palliata*). Folia Primatologica. 1990;54(1-2):1–15.

55. Coleman MN. What do primates hear? a meta-analysis of all known nonhuman primate behavioral audiograms. International Journal of Primatology. 2009;30(1):55–91.

56. Beecher MD. Hearing in the owl monkey (*Aotus trivirgatus*): I. auditory sensitivity. Journal of Comparative and Physiological Psychology. 1974;86(5):898–901.

57. Osmanski MS, Wang X. Measurement of absolute auditory thresholds in the common marmoset (*Callithrix jacchus*). Hearing Research. 2011;277(1):127–133.

58. Beecher MD. Pure-tone thresholds of the squirrel monkey (*Saimiri sciureus*). Journal of the Acoustical Society of America. 1974;55(1):196–198.

59. Heffner RS. Primate hearing from a mammalian perspective. Anatomical Record Part A: Discoveries in Molecular, Cellular, and Evolutionary Biology. 2004;281A(1):1111–1122.

60. Esser KH, Daucher A. Hearing in the FM-bat *Phyllostomus discolor*: a behavioral audiogram. Journal of Comparative Physiology A. 1996;178(6):779–785.

61. Moynihan M. Some behavior patterns of platyrrhine monkeys I. the night monkeys (*Aotus trivirgatus*). Smithsonian Miscellaneous Collection. 1964;146:1–84.

62. Bezerra BM, Souto A. Structure and usage of the vocal repertoire of *Callithrix jacchus*. International Journal of Primatology. 2008;29(3):671–701.

63. Brumm H, Slabbekoorn H. Acoustic communication in noise. Advances in the Study of Behavior. 2005;35:151–209.

64. Waser PM, Brown CH. Is there a “sound window” for primate communication? Behavioral Ecology and Sociobiology. 1984;15(1):73–76.

65. Ellinger N, Hödl W. Habitat acoustics of a neotropical lowland rainforest. Bioacoustics. 2003;13(3):297–321.

66. Fletcher NH. A simple frequency-scaling rule for animal communication. Journal of the Acoustical Society of America. 2004;115(5):2334–2338.

67. Fitch WT. In: Ghazanfar AA, editor. Primate vocal production and its implications for auditory research. Boca Raton: CRC Press; 2003. p. 87–108.

68. Harris TR, Fitch WT, Goldstein LM, Fashing PJ. Black and white colobus monkey (*Colobus guereza*) roars as a source of both honest and exaggerated information about body mass. Ethology. 2006;112(9):911–920.

69. Pope TR. The reproductive consequences of male cooperation in the red howler monkey: paternity exclusion in multi-male and single-male troops using genetic markers. Behavioral Ecology and Sociobiology. 1990;27(6):439–446.

70. Garber PA, Kowalewski MK. In: Sussman RW, Cloninger CR, editors. Collective action and male affiliation in howler monkeys (Alouatta caraya). New York: Springer; 2011. p. 145–165.

71. Fisher RA. The genetical theory of natural selection. Oxford: Clarendon Press; 1930.

72. Ryan MJ, Rand AS. Sexual selection and signal evolution: the ghost of biases past. Philosophical Transactions of the Royal Society of London Series B: Biological Sciences. 1993;340(1292):187–195.

73. Ryan MJ, Kime NM. In: Simmons AM, Popper AN, Fay RR, editors. Selection on long-distance acoustic signals. New York: Springer; 2003. p. 225–274.

74. Searcy WA, Nowicki S. The evolution of animal communication: reliability and deception in signaling systems. Princeton: Princeton University Press; 2005.

75. Fitch WT. Acoustic exaggeration of size in birds via tracheal elongation: comparative and theoretical analyses. Journal of Zoology. 1999;248(1):31–48.

76. Sueur J, Mackie D, Windmill JFC. So small, so loud: extremely high sound pressure level from a pygmy aquatic insect (Corixidae, Micronectinae). PLoS ONE. 2011;6(6):e21089.

77. Bezanson M, Garber PA, Murphy JT, Premo LS. Patterns of subgrouping and spatial affiliation in a community of mantled howling monkeys (*Alouatta palliata*). American Journal of Primatology. 2008;70(3):282–293.

