## Supplemental information for "Audiograms of howling monkeys: are extreme loud calls the result of runaway selection?"

**Figure S1**

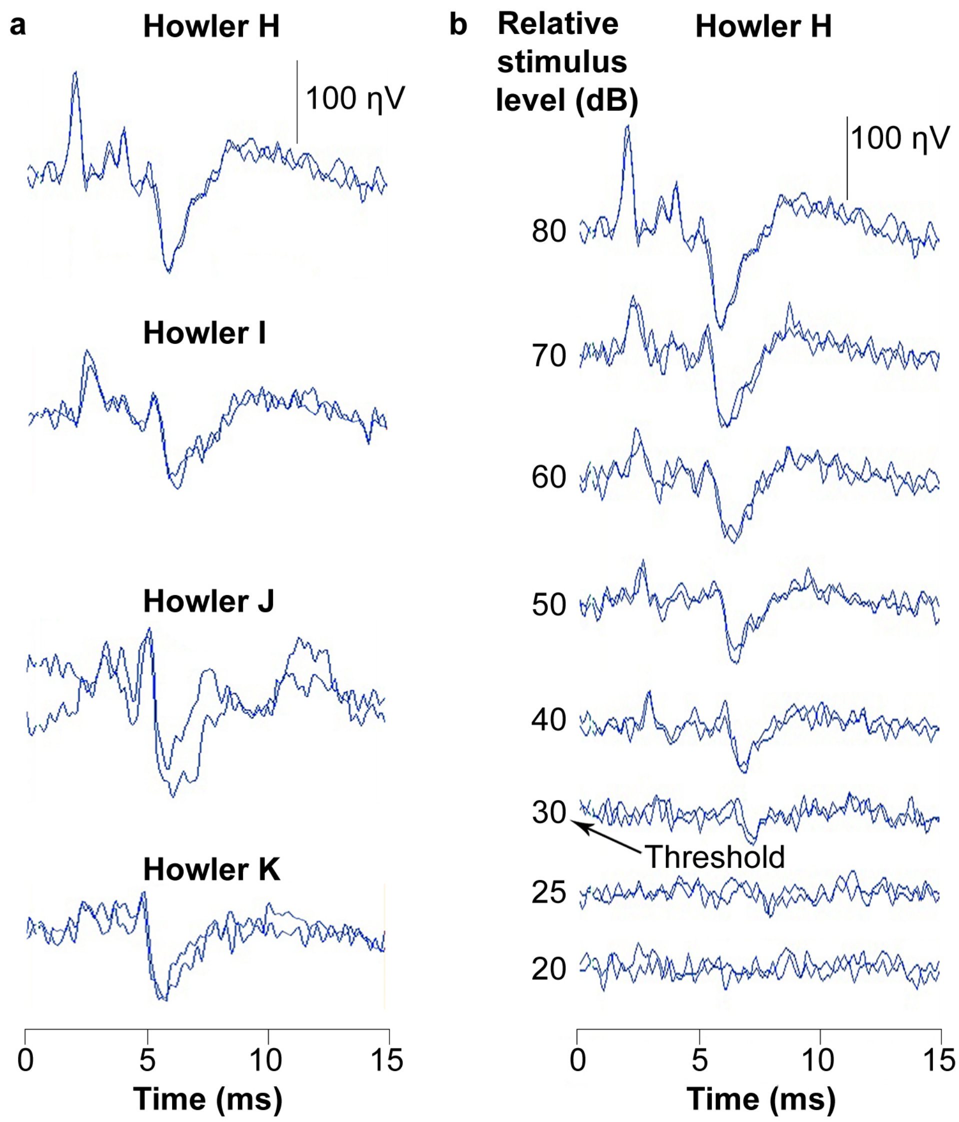


**Figure S1.** Typical ABR waveforms evoked with 11.3 kHz stimuli. A: waveforms for each subject. B: waveform series demonstrating decreasing ABR amplitude with decreasing stimulus level.

**Figure S2**

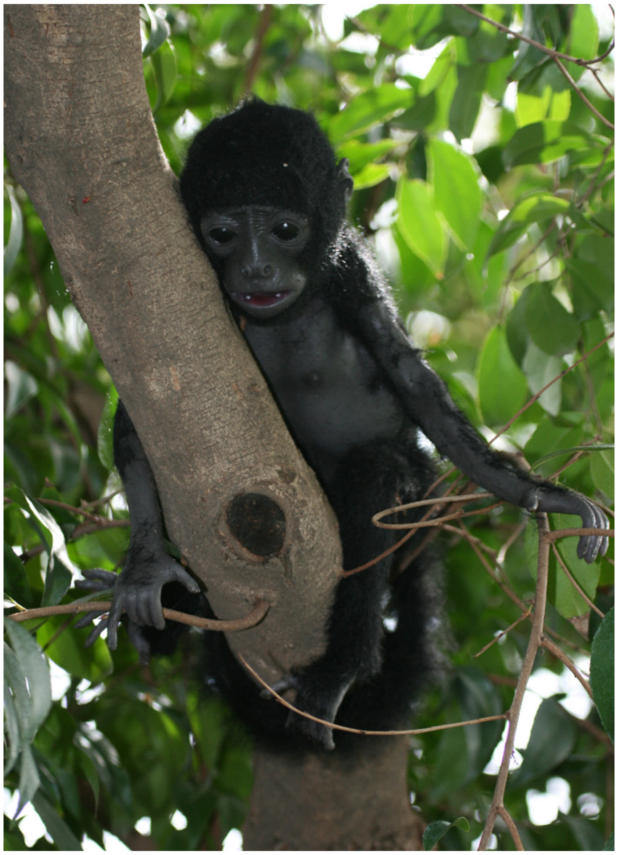

**Figure S2.** Model infant. Field playback experiments of the *wrah-ha* were performed in tandem with a rubber model (Animal Makers Inc., Moorpark, CA, USA). In most conditions, a loudspeaker (Micro Cube, Roland Corp., Hamamatsu, Japan) was positioned next to the model. In one condition, a magnetic speaker (FF1, Tucker-Davis) was embedded in the abdominal cavity of the rubber model. The speaker and model were positioned on tree branches ≈2 m from the ground. On 7 of 11 occasions, the vocalization quickly (within 5 min) attracted lone males or mixed groups. Animals descended within a few meters of the model, but never contacted it. In all cases, adult males and/or females demonstrated heightened vigilance and sustained howling or barking over a span of 5-20 min. On one occasion the call attracted a female black-handed spider monkey (*Ateles geoffroyi*). Such results are equivocal, but they suggest that adults are motivated to approach distressed infants. Our failure to elicit physical interaction with the model could be attributed to the deterrent presence of human observers or the unnatural appearance and vocal 'behavior' of the model. Normative infant behavior predicts the suppression of distress calls in the vicinity of potentially infanticidal males. Another factor was the highly irregular social context. Unweaned infants of *A. palliata* are seldom isolated; they maintain close proximity to their mothers (< 3 m [Clarke, 1990]) and other members of their natal group (1-5 m [Bezanson et al., 2008]). Our model, though lifelike from a distance, may have appeared too unnatural to elicit further response. Photograph by Andrew J. Cunningham.

Supplementary references:

Bezanson, M., Garber, P. A., Murphy, J. T. & Premo, L. S. Patterns of subgrouping and spatial affiliation in a community of mantled howling monkeys (*Alouatta palliata*). *Am. J. Primatol.* **70**, 282-293 (2008).

Clarke, M. R. Behavioral development and socialization of infants in a free-ranging group of howling monkeys (*Alouatta palliata*). *Folia Primatol.* **54**, 1-15 (1990).

**Table S1.** ABR thresholds

| | ABR Threshold (dB SPL) at each frequency (0.5-32 kHz) | | | | | | | | | | | | |
|---|---|---|---|---|---|---|---|---|---|---|---|---|---|
| Subject | 0.5[a,b] | 0.7[a,b] | 1.0[b] | 1.4[b] | 2.0 | 2.8 | 4.0 | 5.7 | 8.0 | 11.3 | 16.0 | 22.6 | 32.0 |
| H | 22.6[c] | 16.7[c] | 15.8[c] | 35.0 | 40.4 | 38.5 | 33.9 | 41.9 | 31.0 | 33.2 | 42.9 | 81.5 | 92.1 |
| I | 54.2 | 44.0 | 35.1 | 37.0 | 41.2 | 49.7 | 51.9 | 56.0 | 33.7 | 33.3 | 44.9 | 72.7 | 87.0 |
| J | 22.5 | 18.7 | 26.3 | 37.9 | 36.0 | 46.8 | 41.7 | 37.5 | 21.4 | 11.9 | 18.3 | 42.7 | 60.8 |
| K | 52.9 | 47.7 | 45.9 | 48.0 | 46.9 | 50.7 | 42.4 | 45.4 | 38.9 | 30.6 | 41.0 | 56.1 | 79.1 |
| Mean | 37.6 | 31.8 | 30.8 | 39.5 | 41.1 | 46.4 | 42.5 | 45.2 | 31.2 | 27.2 | 36.8 | 63.3 | 79.5 |
| St dev | 17.4 | 16.3 | 12.8 | 5.8 | 4.4 | 5.5 | 7.4 | 7.9 | 7.4 | 10.3 | 12.4 | 17.3 | 13.7 |
| Median | 37.5 | 31.3 | 30.7 | 37.4 | 40.8 | 48.2 | 42.0 | 43.7 | 32.4 | 31.9 | 41.9 | 64.4 | 83.1 |
| 1st quart | 22.6 | 18.2 | 23.7 | 36.5 | 39.3 | 44.7 | 39.8 | 40.8 | 28.6 | 25.9 | 35.3 | 52.8 | 74.5 |
| 3rd quart | 52.6 | 44.9 | 37.8 | 40.4 | 42.7 | 49.9 | 44.7 | 48.0 | 35.0 | 33.2 | 43.4 | 74.9 | 88.3 |

[a]Thresholds for 0.5 and 0.7 kHz are preliminary due to ambient noise.
[b]ABR-derived thresholds for frequencies ≤ 2 kHz are generally elevated compared to behavioral-derived thresholds.
[c]Mathematically calculated based on other subjects – see main text for explanation.
